# A comparison of the IgM and IgT repertoires reveals the structure of B cell communities in rainbow trout is a consequence of mechanistic and evolutionary forces

**DOI:** 10.1101/2021.08.18.456864

**Authors:** Gregory R. Costa, Annie Poirier, Erin S. Bromage

## Abstract

In rainbow trout (*Oncorhynchus mykiss*), three classes of antibodies have been identified: IgM, IgD, and IgT, which differ in their abundance and effector functions. Though the process of VDJ recombination that creates these antibodies is often described as a stochastic process, recent findings in mammals suggest that there are biases in antibody construction. Because class switching is absent in teleosts, studying the IgM and IgT communities of rainbow trout provides the opportunity to see how evolution has differentially shaped the IgM and IgT repertoires. Even though it has not yet been demonstrated that biases exist in variable region construction for trout immunoglobulins, it seems reasonable that both natural and artificial selection have driven preferential gene-segment usage and pairing biases in rainbow trout. In this study, we sequenced the heavy chain variable regions of membrane IgM and IgT from multiple fish and predicted that given the more generalist role of the abundant IgM there would be less pressure to bias V gene segment usage and DJ pairings; rather, natural selection would have favored diversity and the ability to respond to a plethora of pathogens. Conversely, as IgT is substantially less abundant in the serum than IgM and specialized in its function, there would be selective pressure to make the most out of a little, thus favoring biased V segment usage and preferential DJ pairings. In support of our hypotheses, for IgT, over 70% of DJ pairs were biased and over 60% of antibodies were constructed with just two V gene segments. These biases were not prevalent in the IgM repertoire, where only 4% of DJ pairs were biased and no single V gene segment was utilized in more than 10% of antibodies. We found that these biases have profound influences in the richness and evenness of the repertoires, with the IgM repertoire investing more equitably in nearly double the number of VDJ combinations compared with IgT.

## Introduction

In rainbow trout, three classes of antibodies have been identified: IgM, IgD, and IgT, which differ in their abundance and effector functions [1]. IgM, whose concentration in the serum is 100-1000x greater than that of IgT, is the primary systemic antibody in fish, while IgT is largely involved in mucosal immunity and microbiota homeostasis. As in mammals, both IgM and IgD are co-expressed and thus somatically recombine using the same V, D, and J segments during antibody construction; however, IgT, which is produced by a distinct subset of B cells from IgM/IgD [1–5], utilizes distinct D and J elements from IgM and IgD while using the same V segments [6, 7].

Though this process of VDJ recombination is often described as a random process, with each D segment having an equal probability of being spliced to a J segment, and each V segment having an equal probability of being spliced to a D segment [8, 9], high-throughput DNA sequencing has recently shown that in humans not all genes are created equal. That is, there is often variation in the frequencies at which V, D, and J segments are utilized. In the human naïve repertoire, for instance, different V genes may be used as little as 0.1% to more than 10% of all rearrangements. [9, 10]. In addition to individual V, D, and J segment biases, it has been demonstrated in mice and in humans that DJ pairings are also biased during the recombination process, with some pairings occurring more frequently and others occurring less frequently than would be expected by chance (i.e., pairings are independent from D and J utilization frequencies) [11–13]. The mechanism that governs combinatorial biases remains unknown. Though the distance between D and J genes certainly influences DJ pairings, it does not adequately explain all biases. An individual’s genetics, recombination signal sequences that flank each V, D, and J genes and accessibility of the segments due to chromatin structure, along with other factors, have all been implicated in biased usage of V, D, and J gene segments and in DJ pairings [9, 12, 14–16]. Such biases have a profound influence an individual’s repertoire. When antibody sequences are obtained from humans, of the thousands of VDJ combinations, the 100 most frequent combinations are responsible for 50% of all recombination events [9, 13]. This discrepancy between actual VDJ frequencies and theoretical frequencies means that it is highly unlikely that an individual will see certain heavy chain/light chain pairings in his or her lifetime.

It would seem paradoxical that, contrary to the need of the immune system to respond to nearly unlimited variations of epitopes, diversity of the repertoire is constrained by biased gene usage and pairings. However, though the immune system can generate antibodies of infinite specificities, *the carrying capacity of B cells is finite*. Hence, in what authors have coined “innate immunological memory [17],” it makes sense that given the limited space, evolutionary pressure of pathogens has driven the production of antibodies directed towards conserved and vulnerable epitopes a pathogen. There is evidence that humans have evolved V-gene-encoded innate immunity to coevolved pathogens, including *Haemophilis influenzae* [18], *Streptococcus pneumoniae* [19], and human cytomegalovirus [17]. Thus, even though it has not yet been demonstrated that biases exist in variable region construction for trout immunoglobulins, it seems reasonable that both natural and artificial selection have driven preferential gene-segment usage and pairing biases in rainbow trout, potentially for resistance to endemic pathogens.

Because class switching is absent in teleosts, studying the IgM and IgT communities of rainbow trout provides the opportunity to see how evolution has differentially shaped the IgM and IgT repertoires. In this study, we sequenced the heavy chain variable regions of membrane IgM and IgT; as IgD is co-expressed with IgM, it was excluded from antibody repertoire analysis. We predicted that given the more generalist role of the abundant IgM there would be less pressure to bias V-gene usage and DJ pairings; rather, natural selection would have favored diversity and the ability to respond to a plethora of pathogens. Consequently, the VDJ repertoire would be more diverse and its structure more even than that of IgT, also reflected in the CDR3 lengths, whose frequency distribution would be symmetrical (i.e., long and short CDR3s tailing from the mean). Conversely, as IgT is substantially less abundant in the serum than IgM and specialized in its function, there would be selective pressure to make the most out of a little. Hence, selection would favor V-gene utilization patterns and DJ combinations of evolutionary protective value. Consequently, the VDJ repertoire would be less diverse and even than that of IgM, with the immune system investing highly in the production of a few VDJ combinations. As a result of the specialty of IgT, natural selection likely would have skewed CDR3 lengths to be either longer or shorter than that of IgM (i.e., the frequency distribution is asymmetrical). With IgM repertoires expected to be more diverse than that of IgT, we then questioned if IgM contained fewer public response among the fish than IgT. With this mindset, we thus explored not only the mechanisms that define the IgM and IgT repertoires but also speculated on the evolutionary pressures for establishing these mechanisms.

## Methods

### Animal maintenance

Rainbow trout were obtained from Blue Streams Hatchery (West Barnstable, MA) and maintained at the University of Rhode Island East Farm Aquaculture Facility (Kingston, RI) and maintained in a 30-gallon tank within a recirculating system using biologically filtered, dechlorinated water. The water was monitored daily for temperature, nitrates, ammonia, and pH. Water temperature was maintained at 6.7 °C, and a light:dark period of 12:12 h was maintained. Fish were fed dry pellet feed (Ziegler Brothers, Gardners, PA) and grown to ~160-200g before the commencement of the study. This study was conducted under the approval of the Institutional Animal Care and Use Committee of the University of Massachusetts Dartmouth and the University of Rhode Island according to the USDA regulations and the Guide for Care and Use of Laboratory Animals.

### Tissue collection and lymphocyte isolation

From eight fish, lymphocytes were harvested from the peripheral blood, kidney (anterior and posterior), and spleen, using Histopaque 1077 (Sigma), as previously described [20]. The cells were resuspended to a concentration of 1×10^7^ cells/ml and stored on ice for same-day use in RNA extraction.

### RNA extraction

Total RNA was extracted from each sample using a combination of TRIzol Reagent (Invitrogen, Carlsbad, CA) and EZ-10 DNAaway RNA Mini-Preps kit (Bio Basic Inc., Markham, ON). Cells were homogenized using glass beads in 1ml TRIzol Reagent and incubated on ice for 30 minutes. To each sample, 200 μl of chloroform was added and mixed vigorously for 60 seconds. The samples were incubated at room temperature for 10 minutes, and centrifuged (12,000 g, 15 min, 4°C). The aqueous phase, containing the extracted RNA, was transferred to BioBasic EZ-10 columns. Total RNA extraction was performed according to the manufacturer’s protocol. Samples were eluted in 30 μl of RNAse-free water, and the RNA purity and concentration were assessed using the NanoDrop (Thermo Fisher Scientific, Waltham, MA). From five of the eight fish, for each immune tissue, equal contributions of RNA were pooled. Thus, RNA from individual fish (fish 1, fish 2, fish 3) and pooled RNA from five fish were used for downstream applications.

### Poly(A) purification, cDNA synthesis, and PCR

A total of 10 μg of total RNA was used for the Poly A^+^ RNA (mRNA) purification of each sample using the Oligotex mRNA Mini Kit (Qiagen, Hilden, Germany) following the manufacturer’s protocol. The first stand cDNA template was synthesized using the 5′ reagents of the Smarter™ RACE cDNA Amplification Kit (Takara USA), following the manufacturer’s protocol; the maximum volume of RNA (10 μl) for each sample was used in the first-strand cDNA synthesis. cDNA was diluted 1:13 and 1:3 in TE for use in PCR targeting the heavy chain of *membrane IgM* and *membrane IgT*, respectively.

IgM reactions were performed using three nested rounds of PCR. Thermocycling was performed under the following conditions: First round, 5 cycles at 94 °C for 30 seconds and 72 °C for 3 minutes; 5 cycles at 94 °C for 30 seconds, 70 °C for 30 seconds, and 72°C for 3 minutes; 20 cycles at 94 °C for 30 seconds, 68°C for 30 seconds, and 72 °C for 3 minutes. Second round, 20 cycles at 94°C, 68°C for 30 seconds, and 72 °C for 3 minutes. Third round, 94°C for 4 minutes; 10 cycles at 94 °C for 30 seconds, 52 °C for 30 seconds, and 72 °C for 35 seconds. The primers used in this study can be found in the supplemental material (S1 Table). All custom oligos were used at a final concentration of 0.2 μM. The commercial 10x universal primer mix (Takara USA) was used at a final 1x concentration. Each PCR reaction was performed in a 50 μl reaction using nuclease-free water to bring to final volume. For the first two rounds, PCR was performed with 2x SeqAMP buffer and SeqAmp DNA polymerase (Takara USA) and for the final amplification, OneTaq 2x Master Mix with Standard Buffer was used (New England BioLabs, Ipswich, MA). As a template in the first round, 2.5 μl of the diluted cDNA was used. The PCR product was diluted (1:50 in TE), and 5 μl was used in the second round PCR. The PCR product of the second round was diluted (1:50 in TE) and 5 μl was used in the final amplification. The primers utilized in the reverse reactions of the first two rounds of PCR were specific to the transmembrane region of IgM, while the final round targeted the Cμ1 region, resulting in an approximately 500-550 bp product containing the entire VDJ exon of IgM. The product was visualized on an agarose gel.

IgT reactions were also performed using three nested steps of PCR. Thermocycling was performed under the following conditions: First and second round, 94 °C for 4 minutes; 30 cycles at 65 °C for 30 seconds, 72 °C for 2 minutes; 72 °C for 4 minutes. Second round, 94 °C for 4 minutes; 30 cycles at 65 °C for 30 seconds, 72 °C for 2 minutes; 72 °C for 4 minutes. Third round, 94 °C for 4 minutes; 10 cycles at 94 °C for 30 seconds, 52 °C for 30 seconds, and 72 °C for 45 seconds. All custom oligos and the commercial universal primer short (Takara USA) were used at a final concentration of 0.2 μM; the commercial 10x universal primer mix (Takara USA) was used at a final 1x concentration. Each PCR reaction was performed in a 50 μl reaction using nuclease-free water to bring to final volume. For the first two rounds, PCR was performed with 2x SeqAMP buffer and SeqAmp DNA polymerase (Takara USA) and for the final amplification, OneTaq 2x Master Mix with Standard Buffer was used (New England BioLabs, Ipswich, MA). As a template in the first round, 2 μl of the diluted cDNA was used. The PCR product was diluted (1:3 in TE), and 5 μl was used in the second round PCR. The PCR product of the second round was diluted (1:50 in TE) and 2 μl was used in the final amplification. The primers utilized in the first two rounds were specific to membrane IgT, with final amplification resulting in an approximately 630 bp product containing the entire VDJ exon of IgT. The product was visualized on an agarose gel.

IgM and IgT PCR products were purified using Agencourt AMPure XP magnetic beads (Beckman Coulter, Brea, CA) according to the manufacturer’s instructions; samples were eluted in 40 μl of TE and concentration determined by NanoDrop (Thermo Fisher) prior to amplicon sequencing.

### Illumina sequencing and pre-processing of Illumina data

The library preparation and sequencing were conducted at the Tufts University Core Sequencing Facility (Boston, MA). Sequencing libraries were prepared using the Nextera DNA XT library prep kit (Illumina, San Diego, CA) and sequenced by 300-bp paired-end MiSeq (Illumina). IgM and IgT repertoire sequencing data were paired in Geneious R8 software package [21](Biomatters, Auckland, New Zealand). As the IgT paired-end reads did not contain sufficient overlap for merging, only IgM paired data were merged, allowing for a mismatch density of 0.1 using the FLASH v1.2.0 plug-in [22]. The merged IgM sequences and paired IgT sequences were converted to FASTA files for submission to IMGT/High V-Quest [23], using program version 3.5.18 (March 11, 2020) and reference directory release 202018-4. The unaligned IgT sequencing submitted to IMGT resulted in two annotations per amplicon: one read of a pair contained most of the V-gene segment and a second read of a pair contained all of the D and J-gene segments (in addition to a small portion of the V-segment). To compensate, for each pair of reads, the IMGT annotation for the high-quality V-segment was combined with the D and J-gene annotations, while the lower-quality V-segment annotation was discarded.

The sequences presented in this chapter have been submitted to NCBI SRA (https://www.ncbi.nlm.nih.gov/sra) under SRA accession PRJNA662346.

### Ecological analyses of the antibody repertoire

Statistical applications typically used in the field of ecology were applied to the IgM and IgT repertoire to both quantify diversity and describe its structure. Analyses were limited to data from the anterior kidney. The IMGT/High V-Quest summary sheet was filtered so that only productive sequences were analyzed. To prevent bias resulting from differences in the sampling depth between IgM and IgT, the total productive reads were subsampled to the minimum number of productive reads for each sample (32,377, 48,467, 45,329, and 52,203 reads for fish 1, fish 2, fish 3, and pooled, respectively). The Shannon’s Equitability Index (*E*_*H*_) and Shannon’s Diversity Index (*H*) were calculated as measures of VDJ diversity and evenness in the IgM and IgT communities. The observed number of unique VDJ combinations was determined, and a conservative estimate of the VDJ diversity was calculated using ACE-1 (abundance-based coverage estimator) of the ChaoSpecies function in the package SpadeR v.0.1.1 [24] in R [25]. Following a test for normality, significant differences in mean diversity and evenness between IgM and IgT were determined with a two-tailed Student’s *t*-test. To visualize the IgM and IgT VDJ community structure, the relative abundance of each VDJ combination was calculated and rank abundance curves were constructed.

The overlap in VDJ utilization and CDR3 sequences of fish was visualized by constructing Venn diagrams using the R package VennDiagram v1.6.20 for both IgM and IgT samples [26]. To prevent bias resulting from differences in the sampling depth between the IgM and IgT samples while preserving as much data as possible, the total productive reads were subsampled to the minimum number of productive reads for each sample (32,377, 48,467, 45,329, and 52,203 reads for fish 1, fish 2, fish 3, and pooled, respectively) before constructing the diagram. We defined a shared response as a VDJ combination or CDR3 sequence shared between at least two fish.

### V_H_ gene segment usage patterns

Patterns of V_H_ genes were visualized by constructing heatmaps of the relative frequencies of V gene segments utilized in productive transcripts of the anterior kidney IgM and IgT. Heatmaps were constructed using the R package gplots v.3.0.4 [27].

To determine if the anterior kidney served as a proxy for the other immune tissues and if utilization patterns were consistent among fish, principal component analysis (PCA) was applied to the relative abundances (percentage of total) of the V-genes utilized for all sampled immune tissues. The PCA was constructed on the scaled dataset using the R package ggplot2 (45) and its extension ggfortify v0.4.8 (46). To determine if there was a statistical difference in V-gene utilization among fish, a Bray-Curtis resemblance matrix was applied to the square-root transformed gene frequencies in Primer 7; the resemblance matrix was then analyzed using an analysis of similarity (ANOSIM) with 999 permutations [28](Auckland, New Zealand).

### Evaluating biases in DJ pairing

To test whether expected pairing frequencies of D and J gene segments was significantly different than the observed frequencies, we used a slightly modified protocol from work by others [12, 13]. In brief, to reduce possible biases arising from the inclusion of multiple sequences from clonal lineages, several precautions were taken: only sequences from the anterior kidney, where clonal expansion in response to antigen should be minimal [20], were analyzed. The data set was filtered to include only nonproductive sequences, which are not under selection due to their inexpression. Next, all apparent clonal lineages were filtered so that only one representative sequence from a clone was included in the final data set. Clones were defined as groups of related sequences that had identical CDR3 sequences and shared V and J genes. The final dataset for each fish contained too few sequences for analysis after filtering, so data from all four samples were merged into a single data set. Next, contingency tables were constructed to test whether pairing frequencies were overrepresented or underrepresented based on overall D and J utilization. Adjusted residuals were used to measure the degree of departure from independence of each pair. For each contingency table, a cell-by-cell test of standardized Pearson residuals was conducted with a significance level of 0.01, adjusted using the Bonferroni method by dividing the significance level by the number of cells in the contingency table. To validate our filtering method, we used a second approach with the algorithm B-cell Repertoire Inductive Lineage and Immunosequence Annotator (BRILIA)[29]. In brief, BRILIA was used to assign sequences to a clonal lineage. The dataset was filtered so that one representative sequence from each clone was included in the final dataset. Statistical analysis was performed as described.

Next, as only IgT displayed notable biases in the nonproductive pairings, only IgT was studied further. To test whether biases in pairings were reflective of expressed antibodies or if subsequent positive or negative selection on B cells skewed DJ frequencies, we also constructed contingency tables for the productive transcripts and compared the results obtained for the nonproductive transcripted (e.g., observed if a particular pairing was either overrepresented or underrepresented in both the nonproductive and productive data sets). Last, DJ recombination in both the productive and nonproductive data sets was visualized with Circos plots by uploading contingency tables to the online tableviewer [30].

### IgM and IgT CDR3 length distributions

Differences in frequency distributions of CDR3 length were compared between IgM and IgT; CDR3 length characteristics were obtained from the immune repertoire pipeline of ARGalaxy [31]. Differences in distributions were tested using a chi-squared test and a difference in mean length between IgM and IgT was tested with a two-tailed Student’s *t*-test.

## Results

### The IgM repertoire is more diverse and even than the IgT repertoire

To understand the VDJ diversity and structure of the antibody repertoire, rank abundance curves were constructed for the IgM and IgT communities for each fish (Fig. 1). It was apparent from the length of these curves that the IgM repertoire contained a greater breadth of VDJ diversity. In the subset data, the IgM repertoire contained 1649-2188 combinations, while the IgT repertoire was nearly half that (Fig. 1; Table 1). The estimated VDJ diversity of 2000-2500 combinations for IgM was significantly greater (p<0.001) than the 1100-1500 combinations observed in IgT (2000-2500). Moreover, as demonstrated by the steep gradient of IgT and the gradual gradient of IgM, the IgM repertoire was more equitable in the abundances of the VDJ combinations (Fig. 1). That is, IgT was largely dominated by a few highly abundant combinations; for instance for every sample, the most abundant VDJ combination contained over 10% of all reads versus approximately 1% for IgM. Not surprisingly, the IgM repertoire was on average more even than the IgT repertoire (*E*_*H*_=0.913±0.00957 and *E*_*H*_ = 0.615±0.115 for IgM and IgT, respectively)(p<0.001)(Table 1). The Shannon Diversity Index (*H*), which takes into account richness and evenness in a community, for the IgM repertoire was on average 1.66x greater than the average for IgT (Table 1).

**Figure 1:**
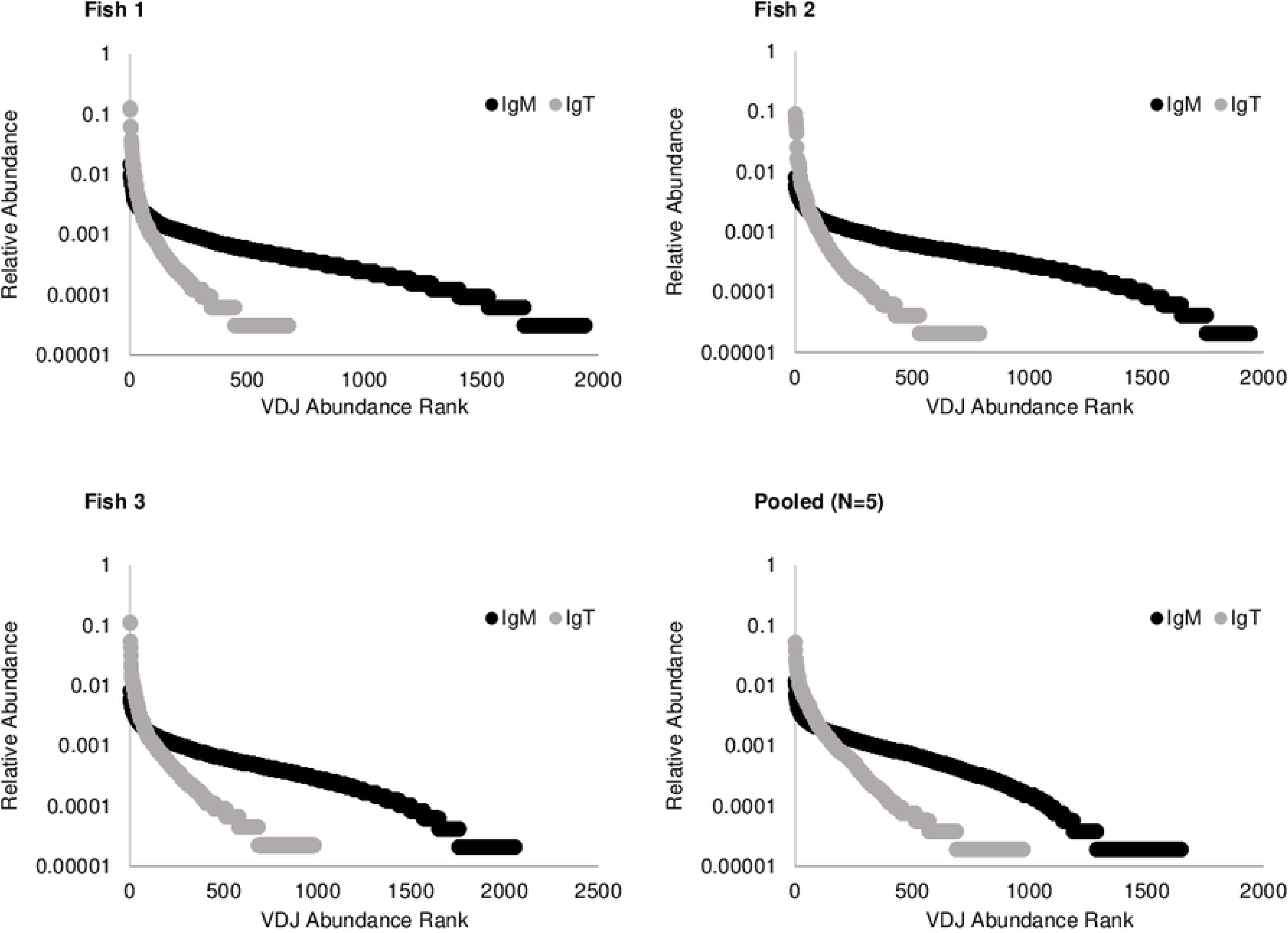
Rank abundance curves for VDJ combinations in IgM and IgT. VDJ abundances are plotted in decreasing order from the most abundant to the least abundant productive VDJ combinations. B cells were harvested from the anterior kidney of three fish and a pooled sample of five fish; the total productive reads for IgM and IgT were subsampled to the minimum number of productive reads for each fish (32,377, 48,467, 45,329, and 53,203 for fish 1, fish 2, fish 3, and pooled, respectively). Note that the relative abundance is scaled logarithmically.

**Table 1:** Diversity indices characterizing the VDJ repertoire of membrane IgM and IgT of four samples.

|  | Fish 1<br>(32,377*) |  | Fish 2<br>(48,467*) |  | Fish 3<br>(45,329*) |  | Pooled<br>(53,200*) |
| --- | --- | --- | --- | --- | --- | --- | --- |
|  | IgM | IgT | IgM | IgT | IgM | IgT | IgM |
| # Unique VDJ Combinations | 1,944 | 678 | 2,056 | 788 | 2,188 | 984 | 1,649 |
| Shannon's Equitability ($E_H$) | 0.91 | 0.61 | 0.92 | 0.63 | 0.92 | 0.47 | 0.90 |
| Shannon Diversity Index ($H$) | 6.85 | 4.00 | 6.99 | 4.22 | 7.05 | 3.23 | 6.68 |
| VDJ Diversity Estimation | 2,116 | 1,103 | 2,346 | 1,265 | 2,392 | 1,503 | 2,478 |

\* Subsampling depth (# productive sequences)

### Biases in antibody construction shape the IgT repertoire

Based on the reduced evenness and diversity present in the IgT repertoire, we then determined the role that V-gene usage and DJ pairing biases play in the production of IgM and IgT. Though IgM and IgT used the same number of V-genes segments (approximately 58), the utilization patters differed substantially, with IgM being diverse and IgT being restricted (Fig. 2). For IgM, though some V segments were used more frequently than others (e.g., V8-30 versus V16-37), no more than 11% of antibodies for a given fish were constructed with a single V gene segment. Rather, IgM construction involved an array of V gene segments often used at similar frequencies. In contrast, the IgT profile was dominated by just a few V gene segments. For all but the pooled sample, 50-60% of all VDJ combinations comprised V1-13 and V9-15. The pooled sample was the most diverse in the utilization patterns, with V1-13 and V9-15 used in about 25% of all antibodies.

**Figure 2:**
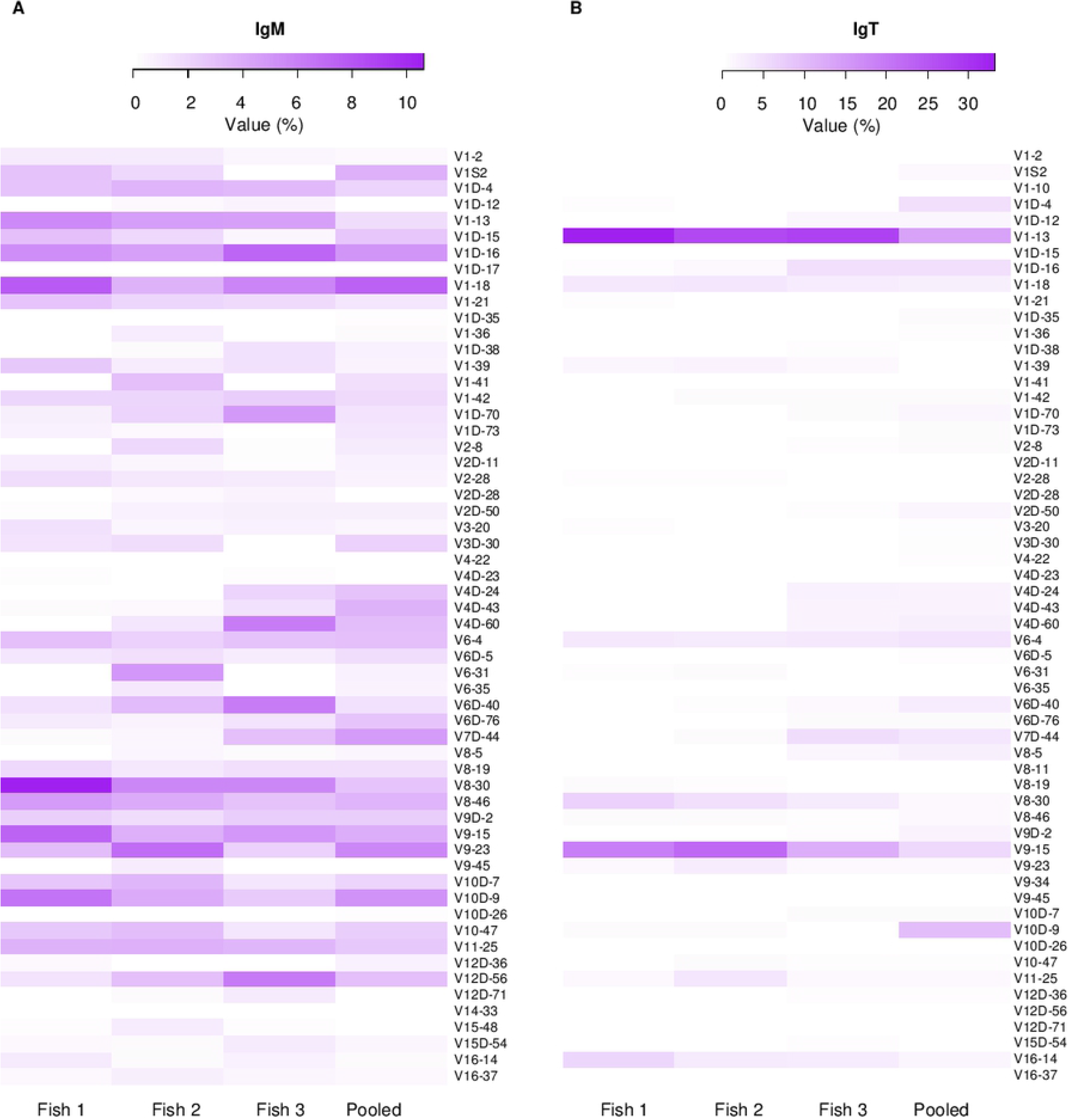
Heatmaps of V_H_ gene usage in (A) IgM and (B) IgT of B cells in the anterior kidney of three fish and a pooled sample comprising five fish. Only productive rearrangements were used to determine the relative frequencies of gene segment usage. Note that (A) and (B) have different color scales.

Several notable findings were revealed from a PCA constructed from the V-gene utilization for the immune tissues from each fish (S1 Fig.). First, usage was related to the fish and not the tissue, as confirmed by an ANOSIM (R=1, p=0.001 for IgM; R=0.962, p=0.001 for IgT). As such the anterior kidney could serve as a proxy for all other tissues in a given fish. Second, while the profiles of IgM were distinct among the fish, in the case of IgT, the confidence intervals for fish 1 and fish 2 overlapped, slightly lowering the R value in the ANOSIM. In addition to differences in V-gene usage, IgM and IgT differ substantially in the number of J-gene segments they utilized and in the frequency distributions. While IgT used four J segments in antibody construction with a single J-gene segment, J1T2, used on average 75% of the time, IgM used six J segments, with no single gene segment being used more than 34% of the time for any given fish (data not shown). The number of D segments and frequency patterns used in VDJ recombination were similar for IgM and IgT.

As recent research in humans and mice uncovered preferential DJ pairings, we asked if these biases would be present in fish and whether they exerted greater influence in shaping the IgT or IgM repertoires. Because the filtering process reduced our statistical power, we were required to combine the data from all four samples. In doing so, we found a substantial difference in the role of preferential pairings in IgM versus IgT: Of the 147 different DJ pairings in nonproductive joins, only four, (2.7%), exhibited any bias in IgM (S2 Table). Two of the pairings occurred more frequently than expected by chance, while two of the pairings occurred less frequently. On the other hand, in IgT, 59 of the 84 DJ nonproductive pairings, or 70%, were biased, and many of these were more extreme than what was seen in IgM (S3 Table; Fig. 3a). For instance, D1T2*01 J2T2 was found 2872 times in our dataset, or three-fold greater than expected. Conversely, D2T1D*01 J2T2 occurred nearly three-times less than expected (S3b Table; Fig. 3a).

**Figure 3:**
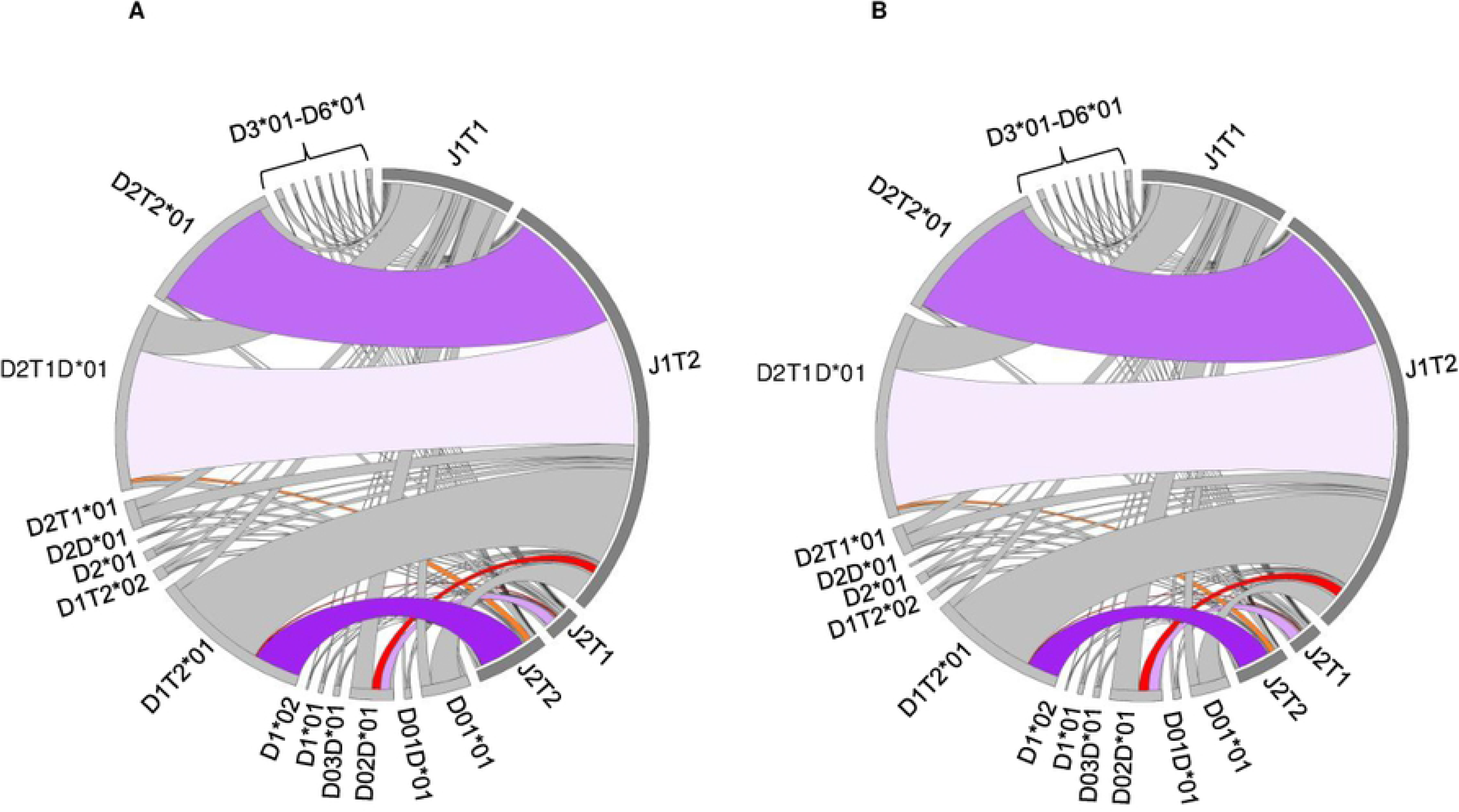
D and J gene associations of (A) nonproductive and (B) productive joins in IgT production reveals biases in pairings. D and J gene associations of **(A)** nonproductive and **(B)** productive joins in IgT production reveals biases in pairings. The sequencing libraries from the anterior kidney four samples were coalesced, and frequency of D-J pairings determined. Links drawn from a D region to a J region indicate an observed DJ recombination, with wider links indicating higher frequencies of observance. For clarity, only 8 highly biased associates of the approximately 59 biased pairings were shaded. Purple shades indicate that the association occurred greater than would be expected at random, while red shades indicate that the association occurred less than would be expected. The darker the shade, the greater the degree of bias. Gray shading indicates pairings with no biases or small but statistically significant biases. A complete list of biased relationships can be seen in S2 Table.

As the biases were minor in the IgM dataset, subsequent analyses were reserved for IgT. We next asked whether biases observed in nonproductive transcripts influenced the functional repertoire, or if positive or negative selection on B cells expressing the resulting antibodies altered these patterns. Indeed, overall, biases present in the nonproductive dataset were reflective of biases present in the productive data set (S3b Table; Fig. 3b). The pattern and frequency of the pairings illustrated in Circos plots were nearly indistinguishable between nonproductive and productive joins (Fig. 3). Similarly, though not all significant biases present in nonproductive rearrangements (S3a Table) were significant in nonproductive rearrangements (S3b Table), the overall trends (i.e., occurring more or less frequently than expected by chance) held true. As tests were conducted with a significance level of 0.01 that was then adjusted with a Bonferroni correction, it is possible additional biases exist that did not exceed the threshold of detection. For instance, had we used an unadjusted value of 0.05, D5T1D*01 J1T1 would have been biased for both nonproductive and productive rearrangements (S3 Table).

### The public response for IgT is not greater than that for IgM

With biases in V and J usage, preferential DJ joins, and fewer J-genes all serving to restrict the IgT repertoire, we then asked if shared responses were more common in IgT than in IgM. Not surprisingly, among the four samples, IgM contained more than double the number of unique VDJ combinations than IgT (3,708 versus 1,749; Fig. 4). However, the degree of sharing was nearly identical between IgM and IgT; the four samples shared 16.4% of all IgM VDJ combinations and 13.7% of all IgT VDJ combinations. In contrast, despite far fewer VDJ combinations utilized in IgT construction, the CDR3 diversity of IgT, with 48530 unique sequences present among the four samples, was more than double than that found in IgM (Fig. 4). This extreme diversity was reflected in the public responses: for IgM, all four samples shared 1% of unique CDR3 amino acid sequences versus 0.5% for IgT. Even when calculations accounted for the abundances of sequences, IgM still displayed a greater degree of sharing. It is important to note that we defined identical CDR3s as having 100% identity. In the case of IgT, different CDR3s sequences for a given combination often differed by a single amino acid (e.g., CAAGGW**G**YAFDYW, CAAGGW**V**YAFDYW, and CAAG**V**WGYAFDYW for the VDJ combination V8-30 D2T1D*01 J1T2), whereas IgM was highly divergent within a given VDJ combination (e.g., CACRTTAAFDYW, CACVCRITARACFDYWO and CAKLYGSGGYFDYW for the VDJ combination V6D-40 D1*01 J2D). Hence, had we been less stringent in how we defined unique CDR3 sequences, the IgM repertoire would likely have been at least as diverse as the IgT repertoire.

**Figure 4:**
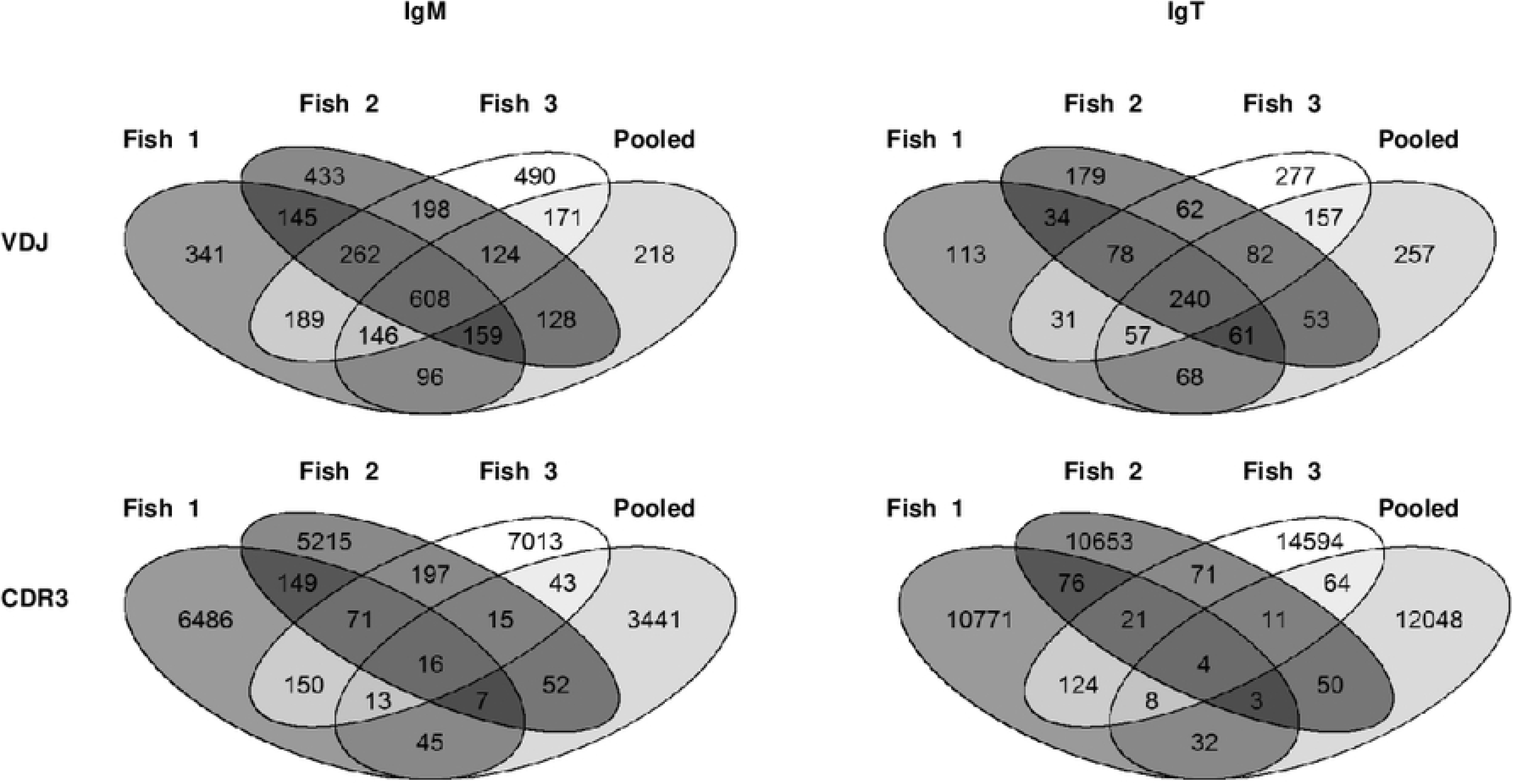
Venn diagrams representing the number of unique IgM/IgT VDJ and CDR3 amino acid sequences that were found to be unique (no overlap) or shared (overlap). IgM and IgT sequencing libraries were obtained from B cells harvested from the anterior kidney of three fish and a pooled sample. So that sampling depth would be equal between IgM and IgT, the total productive reads were subsampled to the minimum number of productive reads for each sample (32377, 48467, 45329, and 53203 for fish 1, fish 2, fish 3, and pooled, respectively).

### The IgT heavy chain is skewed towards longer CDR3s

Last, given that IgT is a specialized antibody largely involved in mucosal immunity, we thought it likely that its CDR3 lengths had been selected to be longer or shorter than those of IgM. The CDR3 lengths of IgM, in contrast, were predicted to exhibit a roughly normal distribution. As predicted, the IgM distribution pattern was a symmetrical one, with a CDR3 length peaking around 10-11 amino acids and lengths tapering off in either direction (Fig. 5). Not only was the distribution skewed in IgT, but also, the frequency distributions for all fish contained two peaks, the largest around 11-12 amino acids, and a second, smaller peak, around 18-20 amino acids. The most striking difference was in the pooled sample; two peaks in IgT, one at 12 amino acids and one at 20 amino acids, were clearly visible. Not surprisingly, the frequency distributions of IgM and IgT for all fish differed significantly when compared with a goodness-of-fit test (p<0.000001, df=18 for all fish). Moreover, the mean IgM CDR3 length of 11.12±0.23 amino acids was significantly shorter than the mean IgT CDR length of 13.09±0.20 amino acids (p=0.00001).

**Figure 5:**
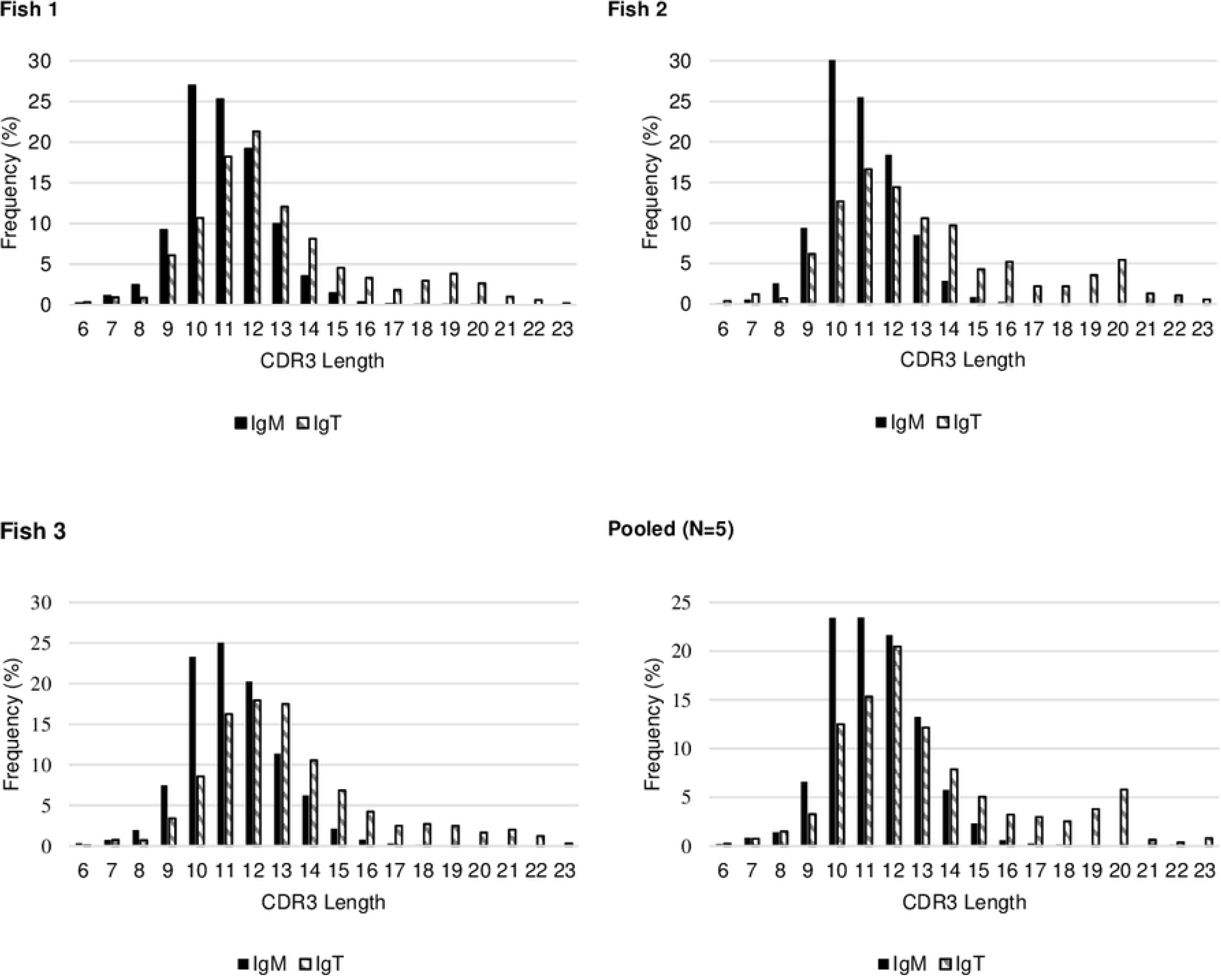
CDR3 length distributions of the anterior kidney IgM and IgT repertoire for the four samples. The distribution profile of CDR3 length was compared by a Chi-square test; for all fish, the profile of IgM significantly differed from that of IgT (p< 0.00001).

## Discussion

Much of the research in the genetics of salmonids is conducted with the goal of ultimately improve breeding programs for traits such as disease resistance [32, 33]. Instead, the main objective of this research was to uncover the mechanisms that shape the antibody repertoire and speculate why these adaptations arose. As teleosts do not class switch their antibodies, rainbow trout provided us with an opportunity unavailable in the mammalian system: Within a single model organism, we were able to compare how evolutionary forces shaped the production of IgM and IgT, which differ significantly in abundance and effector functions [1–4, 34]. We made several predictions about the structure of the respective repertoires and the mechanisms that regulate them. Namely, we predicted that IgT, with its low abundance and specialized function, would contain far fewer VDJ combinations in its repertoire as a consequence of biases, both in V-gene usage and DJ pairing in VDJ recombination. In contrast, IgM, with its broad functions, would be expected to be more diverse to deal with a wide range of pathogens. With a significantly greater carrying capacity for IgM^+^ B cells over IgT^+^ B cells, the benefit of investing in diverse antibodies, some of which likely target rare epitopes, would outweigh the costs. Consequently, biases in antibody construction would be less pronounced. This restriction versus diversity present in IgT versus IgM would thus be reflected in CDR3 lengths, with IgT being selected for possessing either shorter or longer CDR3s than IgM.

Indeed, most of our predictions were substantiated. Not only did IgM contain more than double the number of VDJ combinations in the subset data, but the distribution in frequencies was more even (Table 1; Fig.1). That is, IgT invested more heavily in a few highly abundant VDJ combinations than IgM. Even though these analyses were confined to the anterior kidney, our data suggests that the findings are reflective of utilization patterns of B cells circulating throughout the fish (Fig. S1).

We next asked what causes the discrepancy between the number of combinations in the two isotypes and uncovered that it is an outcome of multiple factors. First, although IgM and IgT used the same number of V segments in antibody production (Fig. 2), IgT invested more than half of antibody production in two to three gene segments (Fig. 2b). The V-gene utilization patterns for IgT were remarkably consistent among all four specimens to the extent that the confidence intervals for two of the fish overlapped in the PCA (Fig. S1); the pooled sample, had, as expected, more diversity than the other three fish but was still far less diverse than what was seen in IgM (Fig. 2). These findings are similar to those observed in torafugu, in which over 50% of IgT antibodies were produced with about three V gene segments, even though IgM and IgT utilized a similar range of V gene segments [7]. This preferential utilization of V segments has also been observed in humans, where abundant alleles may be used 100x more frequently than rarely used alleles. Even similar alleles have been reported to be used at different frequencies; for instance, V1-2*02 is used three times as often as V1-2*04 in individuals who carry both alleles [9, 10]. The findings from torafugu to humans suggest that preferential usage evolved early in the history of vertebrates. Additionally, we found that IgM and IgT differed in the number and investment of utilized J-genes (data not shown). While IgT used four J segments in antibody construction with a single J gene segment used on average 75% of the time, IgM used six J segments, with no single J segment used more than 34% of the time. In torafugu, while IgM used five J segments, the authors found that IgT used a single J segment. Even so, the number of J segments we observed in IgT construction is double the number previously described in rainbow trout [4, 35].

Aside from biases in V and J usage, we asked to what extent DJ-pairing biases exist in IgM and IgT. While under 3% of the DJ pairings of IgM were biased (S2 Table), 70% of the IgT DJ pairings were biased (S3 Table; Fig. 3). Interestingly, researchers have found that in humans, 16 of 186, (8.6%), of observed DJ pairs, exhibited preferential pairing, in contrast to the 27.6% of observed DJ pairs in mice [13]. This suggests that biases in antibody construction developed early in the evolution of the adaptive immune system and became less influential in shaping the repertoire over vertebrate evolution. As in humans [13], we found that the DJ pairings and patterns of biases were comparable for nonproductive and productive rearrangements (S3 Table; Fig. 3), suggesting that the biases are not superseded by positive/negative selection events of B cells and do influence the functional repertoire extensively. There are caveats to our findings: Though the method we used to identify preferential pairings was conservative, we could not control for incorrect annotations of V, D, and J segments in the IMGT database. Over the past two years, the IMGT database has expanded dramatically; if the database is incomplete, we may find that IgT actually contains more D or J gene segments than originally thought. If genes designated as belonging to the same V gene segment actually belong to different segments, then the biases we observed may be an artifact of IMGT’s current limitations. Even though past studies in mammals have found limited evidence of preferential VD segment pairings [12, 13], it would be worthwhile to see if they are present in ancient species. One challenge, as others have encountered, is that the vast number of possible pairs dramatically reduces the statistical power. For us to have been able to test the 1365 potential VD combinations, we would have required a greater sampling depth, as following filtering, the number of reads was reduced by more than half.

Given that IgT had greater biases present than IgM in both segment usage and DJ pairings, we asked if shared responses among the four samples were more common in IgT than IgM. Though the total number of unique IgT VDJ combinations was less than half of the 3700 combinations observed in IgM, the degree of sharing for IgT was nearly identical to IgM (Fig. 4). Surprisingly, the percentage of shared IgT CDR3s was half of what was seen in IgM. However, when we looked at the diversity present in the IgT repertoire, many CDR3s differed by just one amino acid, while the CDR3s in IgM varied by considerably more amino acids. This may represent an adaptation of the two isotypes. Even slight changes in the CDR3 region can dramatically affect antibody binding. Hence, the strategy of IgT-producing B cells may be to create many antibodies that differ slightly in their variable region to target similar epitopes; in doing so, they maximize the chance of creating a high affinity antibody to a narrow range of pathogens. In contrast, IgM must be capable of responding to a broader range of pathogens, and thus a diverse repertoire is constructed with antibodies capable of binding an array of epitopes of many shapes and sizes.

Lastly, given that IgM is both more diverse and less constrained in its antibody construction than IgT, we asked how the frequency distributions of the CDR3 lengths in IgM differed from IgT. We hypothesized that IgT is selected for either larger or shorter CDR3s. The profiles of IgM and IgT were remarkably different; while IgM had a near-symmetrical distribution, the distribution of IgT was reminiscent of directional selection, whereby extreme CDR3 lengths, on average 2 amino acids longer than the IgM length, were favored (Fig. 5). While the profile of IgM consistently had a symmetrical frequency distribution, IgT possessed two peaks: a large peak at approximately 12 amino acids, and a second, smaller peak around 20 amino acids. Again, this likely reflects IgM’s role as a general antibody that must respond to a wide range of threats. In contrast, IgT may have evolved to deal with predictable threats that engage the mucosal system e.g., parasites like *Tetracapsuloides bryosalmonae* and *Enteromyxum leei* where IgT plays the primary role in mucosal protection [35, 36]. Work in torafugu also revealed a narrower IgT CDR3 distribution, albeit the CDR3s were shorter than those observed in IgT in rainbow trout [4]. The shorter CDR3s may be a consequence of the torafugu and rainbow trout immune systems having adapted to their respective environments and thus distinctive pathogen exposure.

In summary, we explored the proximate and ultimate reasons for the structure of the IgM and IgT repertoires. In doing so, we detected some of the mechanisms that restrict diversity, such as preferential V-gene usage, limited usage of J segments, and DJ pairing biases; these mechanisms, pronounced in IgT, are diminished in IgM but are still present. Currently, we do not know how biases and preferential gene usage are implemented but research by others in mice and humans indicates it is complex [9, 12, 14-16]. While we were able to test these proximate mechanisms, we could only speculate on the ultimate reasons. It may be that biases in the IgT repertoire are incidental, with no adaptive value. Alternatively, the forces that shape the IgT repertoire may have nothing to do with immunity but instead in the role of IgT in microbiota homeostasis [5]. We hypothesize, however, that biases are an adaptation to co-evolved pathogens. By limiting the production of IgT to specific VDJ combinations, critical antibodies to common pathogens can be formed, despite being 100-1000x less abundant than IgM. Though this may allow fish to rapidly respond to infectious agents, a downside is that it does not offer the flexibility that class switching provides. This could explain why class switching emerged in amphibians and was retained in subsequent lineages [37]. IgM, on the other hand, has evolved to deal with a plethora of threats, and its repertoire structure more closely resembles that seen in humans (e.g., few biases and thousands of VDJ combinations).

## Supplemental Material

**S1 Table:**
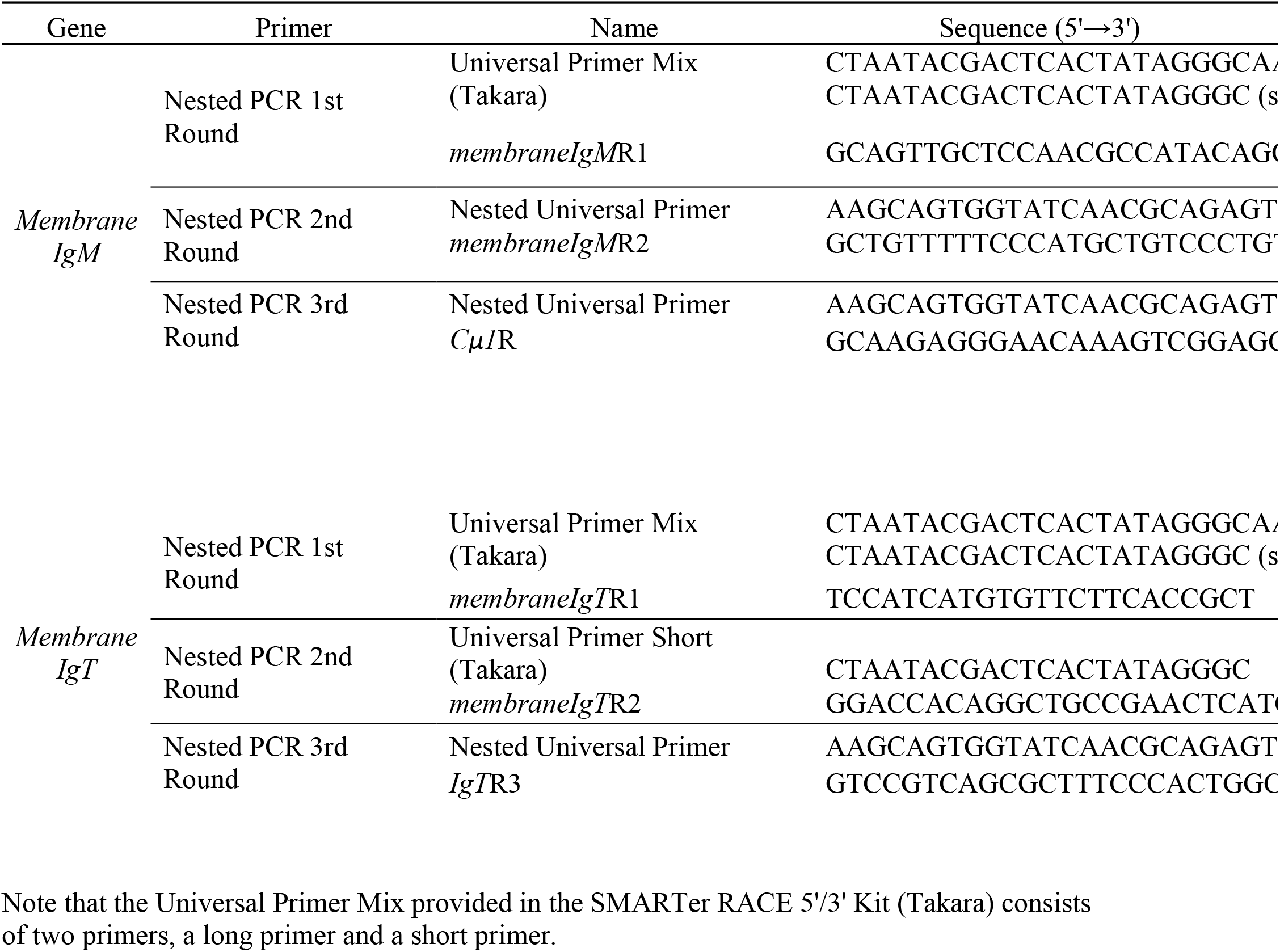
List of oligonucleotide primers used in this study. Primers used in the nested PCR were used to amplify the entire heavy chain variable region of membrane IgM and IgT.

**S2 Table:** Observed versus expected frequencies and adjusted residuals from chi-squared analysis of nonproductive IgM VDJ rearrangement data pooled from four samples.

| VDJ Combination | Observed Frequency | Expected Frequency | Adjusted Residuals |
| --- | --- | --- | --- |
| D1*02 J1T1 | 26 | 2 | 17.59 |
| D1T2*02 J6D | 26 | 5 | 17.59 |
| D3*01 J5 | 12 | 36 | -4.24 |
| D5*01 J5 | 19 | 66 | -6.17 |

Significant biases in D-J pairings are tabulated with their observed and expected frequencies. Adjusted residuals greater than 3.98 (dark gray) represent pairings occurring greater than would be expected by chance, while adjusted residuals less than -3.98 (light gray) represent pairings that occurred less frequently than expected at a 99% confidence interval.

**S1 Figure: Principle component analysis (PCA) applied to V_H_ utilization (A) in IgM production and (B) IgT production in the anterior kidney (AK), peripheral blood (BL), posterior kidney (PK), and spleen (SP).**

Only productive sequences were considered, and the frequencies of all 55 V_H_ genes utilized in IgM and 59 V_H_ genes utilized in IgT were used in PCA calculations.

**S3 Table:** Adjusted residuals from chi-squared analysis of (A) nonproductive and (B) productive IgT rearrangements from data pooled from four samples.

| **A** | J1T1 | J1T2 | J2T1 | J2T2 | **B** | J1T1 | J1T2 | J2T1 | J2T2 |
| --- | --- | --- | --- | --- | --- | --- | --- | --- | --- |
| D01*01 | 5.87 | -2.13 | 6.08 | -8.63 | D01*01 | 8.34 | -4.52 | 5.37 | -9.05 |
| D01D*01 | 10.89 | -11.64 | 7.12 | -1.06 | D01D*01 | 14.93 | -15.50 | 6.45 | -0.50 |
| D02D*01 | 40.23 | -50.86 | 46.14 | -4.36 | D02D*01 | 57.77 | -71.65 | 57.15 | -6.44 |
| D03D*01 | 12.35 | -14.48 | 12.45 | -2.35 | D03D*01 | 13.10 | -19.22 | 21.20 | -2.56 |
| D1*01 | 5.43 | -8.30 | 9.19 | -1.09 | D1*01 | 8.41 | -10.34 | 10.16 | -2.50 |
| D1*02 | 2.95 | -3.92 | 4.14 | -0.79 | D1*02 | 4.68 | -6.91 | 6.06 | 0.35 |
| D1T2*01 | -38.46 | -4.62 | -18.91 | 69.62 | D1T2*01 | -48.19 | -1.84 | -21.72 | 76.36 |
| D1T2*02 | 10.44 | -10.70 | 5.86 | -1.20 | D1T2*02 | 6.94 | -9.57 | 5.31 | 1.81 |
| D2*01 | 10.61 | -14.51 | 11.73 | 0.55 | D2*01 | 11.50 | -18.50 | 15.67 | 3.00 |
| D2D*01 | 6.26 | -4.64 | 3.11 | -2.92 | D2D*01 | 8.07 | -6.83 | 4.13 | -3.54 |
| D2T1*01 | 16.51 | -13.39 | 13.15 | -9.48 | D2T1*01 | 21.48 | -22.79 | 22.11 | -9.93 |
| D2T1D*01 | 13.57 | 14.97 | -16.47 | -28.99 | D2T1D*01 | 19.61 | 10.57 | -24.32 | -30.33 |
| D2T2*01 | -32.22 | 52.13 | -20.75 | -24.85 | D2T2*01 | -50.37 | 69.76 | -27.32 | -24.89 |
| D3*01 | 4.51 | -0.97 | -0.28 | -4.17 | D3*01 | 4.90 | -2.87 | 4.19 | -5.89 |
| D3D*01 | 7.01 | -8.89 | 5.90 | 0.33 | D3D*01 | 12.26 | -12.55 | 5.35 | -0.77 |
| D3T1*01 | 3.73 | -4.97 | 2.00 | 1.21 | D3T1*01 | 3.91 | -3.88 | 2.04 | -0.73 |
| D3T1D*01 | 6.49 | -8.19 | 3.88 | 1.54 | D3T1D*01 | 7.97 | -11.95 | 6.39 | 3.74 |
| D4*01 | 6.93 | -8.60 | 8.17 | -1.43 | D4*01 | 6.96 | -10.71 | 12.57 | -1.48 |
| D5*01 | 3.54 | -3.44 | 2.58 | -0.93 | D5*01 | 1.77 | -3.67 | 3.23 | 1.31 |
| D5T1D*01 | 2.72 | -5.21 | 8.99 | -1.94 | D5T1D*01 | 3.02 | -4.96 | 7.86 | -1.98 |
| D6*01 | 7.24 | -13.64 | 16.24 | 0.50 | D6*01 | 10.98 | -18.30 | 16.62 | 2.69 |

Adjusted residuals greater than 3.85 (dark gray) represent pairings occurring greater than would be expected by chance, while adjusted residuals less than -3.85 (light gray) represent pairings that occurred less frequently than expected at a 99% confidence interval.

## References

1. Hordvik I. Immunoglobulin isotypes in Atlantic salmon, *Salmo salar*. Biomolecules. 2015;5(1):166–77. Epub 2015/03/04. doi: 10.3390/biom5010166. PubMed PMID: 25734583; PubMed Central PMCID: PMCPMC4384117.

2. Zhang YA, Salinas I, Li J, Parra D, Bjork S, Xu Z, et al. IgT, a primitive immunoglobulin class specialized in mucosal immunity. Nat Immunol. 2010;11(9):827–35. Epub 2010/08/03. doi: 10.1038/ni.1913. PubMed PMID: 20676094; PubMed Central PMCID: PMCPMC3459821.

3. Xu Z, Takizawa F, Casadei E, Shibasaki Y, Ding Y, Sauters TJC, et al. Specialization of mucosal immunoglobulins in pathogen control and microbiota homeostasis occurred early in vertebrate evolution. Sci Immunol. 2020;5(44). Epub 2020/02/09. doi: 10.1126/sciimmunol.aay3254. PubMed PMID: 32034088; PubMed Central PMCID: PMCPMC7296778.

4. Castro R, Jouneau L, Pham H-P, Bouchez O, Giudicelli V, Lefranc M-P, et al. Teleost fish mount complex clonal IgM and IgT responses in spleen upon systemic viral infection. PLOS Pathogens. 2013;9(1):e1003098. doi: 10.1371/journal.ppat.1003098.

5. Xu Z, Takizawa F, Casadei E, Shibasaki Y, Ding Y, Sauters TJC, et al. Specialization of mucosal immunoglobulins in pathogen control and microbiota homeostasis occurred early in vertebrate evolution. Science Immunology. 2020;5(44):eaay3254. doi: 10.1126/sciimmunol.aay3254.

6. Zhang YA, Salinas I, Oriol Sunyer J. Recent findings on the structure and function of teleost IgT. Fish Shellfish Immunol. 2011;31(5):627–34. Epub 2011/04/07. doi: 10.1016/j.fsi.2011.03.021. PubMed PMID: 21466854; PubMed Central PMCID: PMCPMC3404837.

7. Fu X, Sun J, Tan E, Shimizu K, Reza MS, Watabe S, et al. High-throughput sequencing of the expressed torafugu (*Takifugu rubripes*) antibody sequences distinguishes IgM and IgT repertoires and reveals evidence of convergent evolution. Front Immunol. 2018;9:251. Epub 2018/03/09. doi: 10.3389/fimmu.2018.00251. PubMed PMID: 29515575; PubMed Central PMCID: PMCPMC5826340.

8. Bonilla FA, Oettgen HC. Adaptive immunity. J Allergy Clin Immunol. 2010;125(2 Suppl 2):S33–40. Epub 2010/01/12. doi: 10.1016/j.jaci.2009.09.017. PubMed PMID: 20061006.

9. Jackson KJ, Kidd MJ, Wang Y, Collins AM. The shape of the lymphocyte receptor repertoire: lessons from the B cell receptor. Front Immunol. 2013;4:263. Epub 2013/09/14. doi: 10.3389/fimmu.2013.00263. PubMed PMID: 24032032; PubMed Central PMCID: PMCPMC3759170.

10. Boyd SD, Gaeta BA, Jackson KJ, Fire AZ, Marshall EL, Merker JD, et al. Individual variation in the germline Ig gene repertoire inferred from variable region gene rearrangements. J Immunol. 2010;184(12):6986–92. Epub 2010/05/25. doi: 10.4049/jimmunol.1000445. PubMed PMID: 20495067; PubMed Central PMCID: PMCPMC4281569.

11. Volpe JM, Kepler TB. Large-scale analysis of human heavy chain V(D)J recombination patterns. Immunome Res. 2008;4:3. Epub 2008/02/29. doi: 10.1186/1745-7580-4-3. PubMed PMID: 18304322; PubMed Central PMCID: PMCPMC2275228.

12. Kidd MJ, Jackson KJ, Boyd SD, Collins AM. DJ pairing during VDJ recombination shows positional biases that vary among individuals with differing IGHD locus immunogenotypes. J Immunol. 2016;196(3):1158–64. Epub 2015/12/25. doi: 10.4049/jimmunol.1501401. PubMed PMID: 26700767; PubMed Central PMCID: PMCPMC4724508.

13. Arnaout R, Lee W, Cahill P, Honan T, Sparrow T, Weiand M, et al. High-resolution description of antibody heavy-chain repertoires in humans. PLoS One. 2011;6(8):e22365. Epub 2011/08/11. doi: 10.1371/journal.pone.0022365. PubMed PMID: 21829618; PubMed Central PMCID: PMCPMC3150326.

14. Greiff V, Menzel U, Miho E, Weber C, Riedel R, Cook S, et al. Systems analysis reveals high genetic and antigen-driven predetermination of antibody repertoires throughout B cell development. Cell Rep. 2017;19(7):1467–78. Epub 2017/05/18. doi: 10.1016/j.celrep.2017.04.054. PubMed PMID: 28514665.

15. Rubelt F, Bolen CR, McGuire HM, Vander Heiden JA, Gadala-Maria D, Levin M, et al. Individual heritable differences result in unique cell lymphocyte receptor repertoires of naive and antigen-experienced cells. Nat Commun. 2016;7:11112. Epub 2016/03/24. doi: 10.1038/ncomms11112. PubMed PMID: 27005435; PubMed Central PMCID: PMCPMC5191574.

16. Wang C, Liu Y, Cavanagh MM, Le Saux S, Qi Q, Roskin KM, et al. B-cell repertoire responses to varicella-zoster vaccination in human identical twins. Proc Natl Acad Sci U S A. 2015;112(2):500–5. Epub 2014/12/24. doi: 10.1073/pnas.1415875112. PubMed PMID: 25535378; PubMed Central PMCID: PMCPMC4299233.

17. Thomson CA, Bryson S, McLean GR, Creagh AL, Pai EF, Schrader JW. Germline V-genes sculpt the binding site of a family of antibodies neutralizing human cytomegalovirus. EMBO J. 2008;27(19):2592–602. Epub 2008/09/06. doi: 10.1038/emboj.2008.179. PubMed PMID: 18772881; PubMed Central PMCID: PMCPMC2567409.

18. Lucas AH, Reason DC. Polysaccharide vaccines as probes of antibody repertoires in man. Immunological Reviews. 1999;171(1):89–104. doi: 10.1111/j.1600-065X.1999.tb01343.x.

19. Zhou J, Lottenbach KR, Barenkamp SJ, Lucas AH, Reason DC. Recurrent variable region gene usage and somatic mutation in the human antibody response to the capsular polysaccharide of *Streptococcus pneumoniae* type 23F. Infect Immun. 2002;70(8):4083–91. Epub 2002/07/16. doi: 10.1128/iai.70.8.4083-4091.2002. PubMed PMID: 12117915; PubMed Central PMCID: PMCPMC128163.

20. Bromage ES, Kaattari IM, Zwollo P, Kaattari SL. Plasmablast and plasma cell production and distribution in trout immune tissues. J Immunol. 2004;173(12):7317–23. Epub 2004/12/09. doi: 10.4049/jimmunol.173.12.7317. PubMed PMID: 15585855.

21. Kearse M, Moir R, Wilson A, Stones-Havas S, Cheung M, Sturrock S, et al. Geneious basic: an integrated and extendable desktop software platform for the organization and analysis of sequence data. Bioinformatics. 2012;28(12):1647–9. Epub 2012/05/01. doi: 10.1093/bioinformatics/bts199. PubMed PMID: 22543367; PubMed Central PMCID: PMCPMC3371832.

22. Magoc T, Salzberg SL. FLASH: fast length adjustment of short reads to improve genome assemblies. Bioinformatics. 2011;27(21):2957–63. Epub 2011/09/10. doi: 10.1093/bioinformatics/btr507. PubMed PMID: 21903629; PubMed Central PMCID: PMCPMC3198573.

23. Alamyar E, Duroux P, Lefranc MP, Giudicelli V. IMGT((R)) tools for the nucleotide analysis of immunoglobulin (IG) and T cell receptor (TR) V-(D)-J repertoires, polymorphisms, and IG mutations: IMGT/V-QUEST and IMGT/HighV-QUEST for NGS. Methods Mol Biol. 2012;882:569–604. Epub 2012/06/06. doi: 10.1007/978-1-61779-842-9_32. PubMed PMID: 22665256.

24. Chao A, Ma KH, Hsieh TC. SpadeR: species prediction and diversity estimation with R. R package version 0.1.0. Retrieved from http://chao.stat.nthu.edu.tw/blog/software-download/2015.

25. R Development Core Team. R: a language and environment for statistical computing. Retrieved from http://www.R-project.org. Vienna, Austria: R Foundation for Statistical Computer; 2010.

26. Chen H, Boutros PC. VennDiagram: a package for the generation of highly-customizable Venn and Euler diagrams in R. BMC Bioinformatics. 2011;12:35. Epub 2011/01/29. doi: 10.1186/1471-2105-12-35. PubMed PMID: 21269502; PubMed Central PMCID: PMCPMC3041657.

27. Warnes G, Bolker B, Bonebakker L, Gentleman R, Liaw W, Lumley T, et al., editors. gplots: various R programming tools for plotting data. R package version 3.0.4. Retrieved from https://cran.r-project.org/web/packages/gplots/gplots.pdf2015.

28. Anderson M, Gorley R, Clarke K, Anderson M, Gorley R, Clarke K, et al. PERMANOVA+ for PRIMER. Guide to software and statistical methods. 2008.

29. Lee DW, Khavrutskii IV, Wallqvist A, Bavari S, Cooper CL, Chaudhury S. BRILIA: integrated tool for high-throughput annotation and lineage tree assembly of B-cell repertoires. Front Immunol. 2016;7:681. Epub 2017/02/02. doi: 10.3389/fimmu.2016.00681. PubMed PMID: 28144239; PubMed Central PMCID: PMCPMC5239784.

30. Krzywinski MI, Schein JE, Birol I, Connors J, Gascoyne R, Horsman D, et al. Circos: an information aesthetic for comparative genomics. Genome Research. 2009. doi: 10.1101/gr.092759.109.

31. IJspeert H, van Schouwenburg PA, van Zessen D, Pico-Knijnenburg I, Stubbs AP, van der Burg M. Antigen Receptor Galaxy: a user-friendly, web-based tool for analysis and visualization of T and B cell receptor repertoire data. J Immunol. 2017;198(10):4156–65. Epub 2017/04/19. doi: 10.4049/jimmunol.1601921. PubMed PMID: 28416602; PubMed Central PMCID: PMCPMC5421304.

32. Lhorente JP, Araneda ME, Neira R, Yáñez JM. Advances in genetic improvement for salmon and trout aquaculture: the Chilean situation and prospects. Reviews in Aquaculture. 2019;11:340–53.

33. Yanez JM, Houston RD, Newman S. Genetics and genomics of disease resistance in salmonid species. Front Genet. 2014;5:415. Epub 2014/12/17. doi: 10.3389/fgene.2014.00415. PubMed PMID: 25505486; PubMed Central PMCID: PMCPMC4245001.

34. Dong F, Yin G-m, Meng K-f, Xu H-y, Liu X, Wang Q-c, et al. IgT plays a predominant role in the antibacterial immunity of rainbow trout olfactory organs. Frontiers in Immunology. 2020;11(2942). doi: 10.3389/fimmu.2020.583740.

35. Abos B, Estensoro I, Perdiguero P, Faber M, Hu Y, Díaz Rosales P, et al. Dysregulation of B cell activity during proliferative kidney disease in rainbow trout. Frontiers in Immunology. 2018;9(1203). doi: 10.3389/fimmu.2018.01203.

36. Piazzon MC, Galindo-Villegas J, Pereiro P, Estensoro I, Calduch-Giner JA, Gómez-Casado E, et al. Differential modulation of IgT and IgM upon parasitic, bacterial, viral, and dietary challenges in a perciform fish. Frontiers in Immunology. 2016;7(637). doi: 10.3389/fimmu.2016.00637.

37. Van Epps HL. Evolution of class switching. J Exp Med. 2005;202(6):724–. doi: 10.1084/jem2026iti3. PubMed PMID: PMC2212931.

